# Outdoor Air Pollution Relates to Amygdala Subregion Volume and Apportionment in Early Adolescents

**DOI:** 10.1101/2024.10.14.617429

**Authors:** Jessica Morrel, L. Nate Overholtzer, Kirthana Sukumaran, Devyn L. Cotter, Carlos Cardenas-Iniguez, J. Michael Tyszka, Joel Schwartz, Daniel A. Hackman, Jiu-Chiuan Chen, Megan M. Herting

## Abstract

**Background:** Outdoor air pollution is associated with an increased risk for psychopathology. Although the neural mechanisms remain unclear, air pollutants may impact mental health by altering limbic brain regions, such as the amygdala. Here, we examine the association between ambient air pollution exposure and amygdala subregion volumes in 9–10-year-olds.

**Methods:** Cross-sectional Adolescent Brain Cognitive Development^SM^ (ABCD) Study^®^ data from 4,473 participants (55.4% male) were leveraged. Air pollution was estimated for each participant’s primary residential address. Using the probabilistic CIT168 atlas, we quantified total amygdala and 9 distinct subregion volumes from T1- and T2-weighted images. First, we examined how criteria pollutants (i.e., fine particulate matter [PM_2.5_], nitrogen dioxide, ground-level ozone) and 15 PM_2.5_ components related with total amygdala volumes using linear mixed-effect (LME) regression. Next, partial least squares correlation (PLSC) analyses were implemented to identify relationships between co-exposure to criteria pollutants as well as PM_2.5_ components and amygdala subregion volumes. We also conducted complementary analyses to assess subregion apportionment using amygdala relative volume fractions (RVFs).

**Results:** No significant associations were detected between pollutants and total amygdala volumes. Using PLSC, one latent dimension (LD) (52% variance explained) captured a positive association between calcium and several basolateral subregions. LDs were also identified for amygdala RVFs (ranging from 30% to 82% variance explained), with PM_2.5_ and component co-exposure associated with increases in lateral, but decreases in medial and central, RVFs.

**Conclusions:** Fine particulate and its components are linked with distinct amygdala differences, potentially playing a role in risk for adolescent mental health problems.

**Graphical Abstract:** 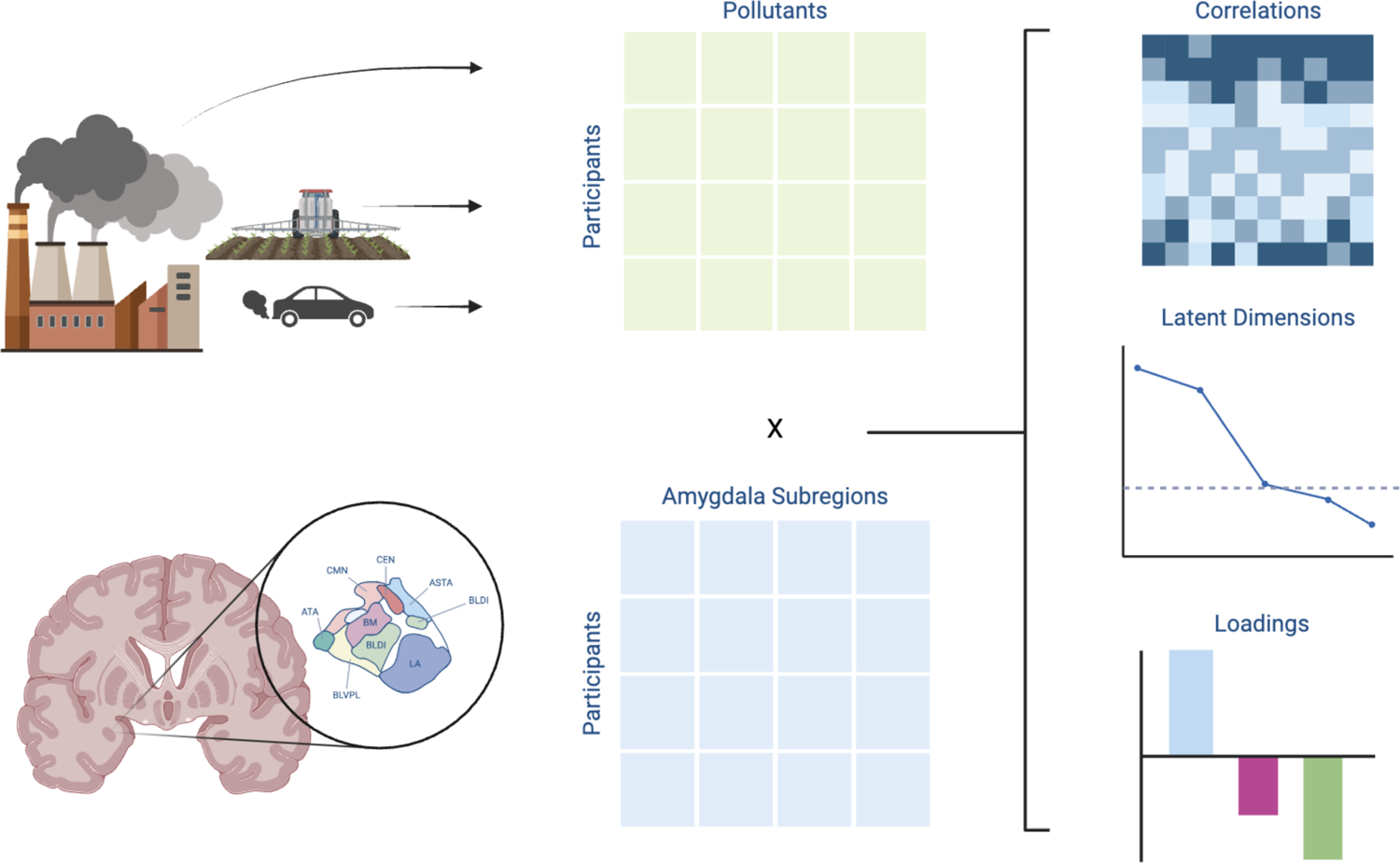

## 1. Introduction

Nearly one in five children in the United States suffer from mental health conditions (1). The average age of onset for these concerns is adolescence, peaking at 14.5 years of age (2). While many social, environmental, and genetic risk factors have been identified, the specific etiology of most psychopathologies remains largely unknown (3). Understanding the risk factors potentially associated with the emergence of these conditions is critical for prevention and early intervention. In this regard, mounting evidence supports a relationship between outdoor air pollution exposure and risk for psychopathology during childhood and adolescence, albeit with mixed findings depending on pollutants, exposure timings, and outcomes (4–7). Ambient air pollution is a ubiquitous mixture of chemicals, increasing recognized as a pervasive neurotoxicant (8,9), and associated with many neurodevelopmental (10–12) and mental health (13–15) conditions. Several criteria pollutants are routinely monitored, including ground-level ozone (O_3_), nitrogen dioxide (NO_2_), and particulate matter (PM) (16). O_3_ is produced through chemical reactions of nitrogen oxides and volatile organic compounds (17), whereas NO_2_ is emitted by vehicles and power plants when fuel is burned (18). PM is composed of many particles and liquid droplets through chemical reactions between pollutants and other sources (19). Fine particulate matter (PM_2.5_; <2.5μm) and smaller particles are emerging as potentially the most harmful of all common pollutants for human health, due to their ability to penetrate deeper into the lung and cross into the bloodstream (20,21). The components in PM_2.5_ can be further classified, and have different environmental origins and chemical features (22). Studying criteria air pollutants along with PM components allows for a more robust understanding of the contributions of different pollution sources and the mental health implications of distinct pollutants (23).

The brain mechanisms linking air pollution and risk for psychopathologies remain unclear. Inhaled pollutants are translocated to the nasal cavity and lungs, where they can cross into the bloodstream and circulate throughout the body (24), causing peripheral inflammation and oxidative stress (25). Some particles can directly access the brain by crossing the blood-brain barrier (BBB), which can induce BBB and neuronal damage, cell death, and a cascade of neuroinflammation (26). Emerging evidence also suggests that inhalation of metals plays a key role in neurotoxicity of airborne particles, causing neuronal cell death, oxidative stress, and inducing dyshomeostasis in the brain (27,28). Among PM components, heavy metals and organic compounds are emerging as particularly toxic for human health (29–31). In addition to these cellular and molecular processes, the amygdala is one brain region by which air pollution may affect mental health. The amygdala is a subcortical, limbic structure that plays a role in emotion regulation, fear, and social behavior (32). Early adolescence marks a critical period of amygdala development, as it reaches its peak total volume at 9-11 years (33), while the development of cortico-limbic circuitry continues through adolescence (34,35). While neuroimaging research has typically investigated the amygdala as a singular structure, it is composed of functionally and cytoarchitecturally distinct subregions (36), which have been associated with emotional processing (37), anxiety (39,43,44), and neurodevelopmental (38,39) and mood (40–42) disorders. However, studies examining associations between air pollution and amygdala structure during childhood and adolescence are few, with mixed findings. While three studies have linked prenatal air pollution exposure with amygdala volumes in infants (45) and 9-12 year-olds (46,47), several others have failed to detect a relationship between prenatal or childhood/adolescent air pollution exposure and amygdala volumes in children/adolescents (46,48–50). Furthermore, no study has assessed the relationship between air pollution exposure and amygdala subregions, nor have they explored the impact of co-exposures to PM_2.5_ components on amygdala subregion volumes.

The aim of the current study was to fill these gaps by examining the relationships between ambient air pollutants and amygdala subregion volumes in 9–10-year-olds. Exposures to annual criteria pollutants and PM_2.5_ components were estimated for each participant’s primary residential address at the time of their MRI scan. We implemented the high-resolution, in vivo probabilistic CIT168 atlas to bilaterally segment total amygdala volumes and nine amygdala subregions for each participant (51,52). We then conducted a series of analyses to characterize the associations between air pollution and amygdala volumes. First, we implemented linear mixed-effect (LME) models to investigate whether each pollutant independently related to total amygdala volume. Next, considering that individuals are not exposed to a given pollutant in isolation (53), we implemented a multivariate data-driven method, partial least squares correlation (PLSC) to identify associations between co-exposure to pollutants and amygdala outcomes. We ran four PLSC analyses examining relationships between two classes of air pollutants, criteria air pollutants and PM_2.5_ components, and nine amygdala subregions of interest. We conducted these analyses first using probabilistic subregion volumes. Then, to account for differences in the relative apportionment of the subregions, we ran a second set of analyses using the relative proportion of each subregion volume to total hemispheric amygdala volume, hereby referred to as amygdala relative volume fractions (RVFs). Given the number of studies failing to find an association between air quality and amygdala volumes, we hypothesized that air pollutants may not be associated with total amygdala volume, but rather co-exposure to PM_2.5_ components would be associated with distinct amygdala subregion differences at 9-10 years old.

## 2. Methods and Materials

### 2.1 Study Design

The current study utilized a subsample of cross-sectional data collected on Siemens 3T MRI scanners from the ongoing Adolescent Brain Cognitive Development^SM^ (ABCD) Study^®^ obtained during the baseline enrollment visit (NIMH Data Archive [NDA] annual 3.0 [imaging data] and 5.0 [all other data] releases; 44,45) The ABCD Study implemented identical recruitment protocols to enroll 11,880 9- and 10-year-old children (mean age = 9.49; 48% female) from 21 sites between October 2016 and October 2018 across the United States in a 10-year longitudinal study (56–58). Centralized Institutional Review Board (IRB) approval was obtained from the University of California San Diego, and each of the 21 study sites obtained approval for experimental and consent procedures from their local IRB. Each child’s parent or legal guardian provided written consent for their child to participate in the study; each child also provided their written assent. Participants were eligible for enrollment in the ABCD Study if they were 9.0-10.99 years at the baseline visit and were fluent in English. Participants were excluded if they had severe sensory, neurological, medical, or intellectual limitations, or if they were unable to complete the MRI scan. A thorough description of study design and procedures can be found elsewhere (56,58).

### 2.2 Participant Sample Selection

A total of 4,473 participants were included in the current analytic sample (**Table 1**, **Figure S1**). Given the CIT168 atlas was developed and validated using T1-weighted (T1w) and T2-weighted (T2w) imaging data collected on a Siemens MRI scanner (51,52) and large differences have been reported in the ABCD study due to scanner manufacturer (59,60), 7,273 participants with neuroimaging data collected on Siemens MRI scanners from 13 ABCD Study sites were eligible for the current study (**Table S1**). Of these, 6,525 met quality control criteria set by the ABCD Study for T1w and T2w image inclusion and 6,449 successfully passed amygdala segmentation by our team. To ensure estimates were reliable within individual amygdalae, we also required each participant to have an intra-amygdala contrast-to-noise ratio (CNR) > 1.0 for T1w and T2w images, as previously published in detail for this technique (see 49 supplemental data). Of the preprocessed participants, 4,754 participants had a CNR > 1.0 in either hemisphere of T1w and T2w images and were therefore considered to have high-quality amygdala segmentations (52,61,62). Due to the nature of PLSC, we were only able to include participants with complete listwise data (i.e., exposures, covariates, and brain data). From the 4,754 participants with usable amygdala data, 281 participants were removed due to missingness (see **Table S2** for comparison between analytic and whole ABCD samples).

**Table 1.** Demographics for analytic sample. . Values represent N (frequency) or Mean (Standard Deviation). The parent report Neighborhood Safety/Crime Survey (NSC) is on a 1-5 scale with 5 indicating a higher degree of perceived neighborhood safety. Note: Race/Ethnicity categories reported in table were collapsed into smaller categories in final analytic models; see **Table S2** for categories used in analyses.

|  |  |
| --- | --- |
| **N total participants** | 4,473 |
| **Age (years)** | 9.98 (0.63) |
| **Sex**<br>Female<br>Male | 1996 (44.6%)<br>2477 (55.4%) |
| **Race/Ethnicity**<br>Asian<br>American Indian / Native American / Native Hawaiian /<br>Other Pacific Islander / Other<br>Hispanic<br>Non-Hispanic Black<br>Non-Hispanic White<br>Multiracial<br>Missing/Refused<br>Don't Know | 63 (1.3%)<br>34 (0.8%)<br>807 (18.0%)<br>586 (13.1%)<br>2622 (58.6%)<br>340 (7.6%)<br>19 (0.4%)<br>2 |
| **Household Income**<br>< \$50k<br>≥ \$50k & < \$100k<br>≥ \$100k<br>Don't Know/Refused | 1067 (23.9%)<br>1241 (27.7%)<br>1847 (41.3%)<br>318 (7.1%) |
| **Highest Parental Education**<br>< HS Diploma<br>HS Diploma/General Educational Development<br>Some College/Associate's Degree<br>Bachelor's degree<br>Post-Graduate Degree<br>Missing/Refused | 129 (2.9%)<br>390 (8.7%)<br>1111 (24.8%)<br>1256 (28.1%)<br>1581 (35.3%)<br>6 (0.1%) |
| **Urbanicity**<br>Rural Area<br>Urban Clusters<br>Urbanized Area | 335 (7.5%)<br>156 (3.5%)<br>3982 (89.0%) |
| **BMI Z-Score** | 0.3 (1.2) |
| **Weekly Physical Activity (days)** | 3.6 (2.3) |
| **Avg. Daily Screen Time (hours)** | 3.0 (2.4) |
| **Neighborhood Safety** | 3.9 (0.9) |

### 2.3 Air Pollution Exposure Estimates

Annual average concentrations of ambient air pollution exposure were estimated at the addresses of each child, as previously described (63). In brief, daily estimates of PM_2.5_ (µg/m^3^) and NO_2_ (ppb), and daily 8-hour maximums of O_3_ (ppb) were derived at a 1-km^2^ resolution using hybrid spatiotemporal models, which utilize satellite-based aerosol optical depth models, land-use regression, weather data, and chemical transport models (64–66). Similarly, annual mean concentrations to fifteen PM_2.5_ components—bromine (Br), calcium (Ca), copper (Cu), elemental carbon (EC), iron (Fe), potassium (K), ammonium (NH_4_^+^), nickel (Ni), nitrate (NO_3_^-^), organic carbon (OC), lead (Pb), silicon (Si), sulfate (SO_4_^2-^), vanadium (V), zinc (Zn)—were estimated monthly at a 50-meter spatial resolution using hybrid spatiotemporal models as previously described (67,68). These exposure estimates were then averaged for the 2016 calendar year to correspond to the baseline enrollment period of the ABCD Study and assigned to the primary residential address of each child provided by the caregiver at the baseline study visit.

### 2.4 Neuroimaging Data

As previously published, a harmonized data protocol was utilized across all ABCD study sites (69). Motion compliance training and real-time, prospective motion correction were used to reduce motion distortion. T1w images were acquired using a magnetization-prepared rapid acquisition gradient echo sequence and T2w images were obtained with a fast spin echo sequence with variable flip angle as previously described (69). Both acquisitions consist of 176 slices with 1 mm^3^ isotropic resolution. T1w and T2w images were then reviewed by ABCD study staff; only data meeting quality control standards for recommended inclusion and those without clinical findings (70) were included. Raw T1w and T2w images were then downloaded from the ABCD 3.0 release (NDA 3.0 data release 2023; 54).

Details of the CIT168 atlas construction, validation, comparison with other atlases, and individual difference estimates are described elsewhere (51,52). Detailed descriptions of each subregion from this atlas along with its successful application to estimate amygdala subregions in children and adolescents have been previously published (61,62,71,72). For each participant, we quantified in vivo probabilistic volumes for nine amygdala subregions of interest per hemisphere as previously described (61,73). Briefly, prior to segmentation, T1w and T2w images were registered via the Human Connectome Project minimal preprocessing pipeline (74). Next, we utilized the CIT168 atlas, a high-resolution, in vivo probabilistic atlas of human amygdala nuclear subregions, to segment each individual’s amygdala into nine subregions (51,52). B-spline bivariate symmetric normalization diffeomorphic registration algorithm from ANTs version 2.2.0 was adapted for image registration of T1w and T2w images to the CIT168 atlas (**Figure 1A**) (75). The inverse diffeomorphism was then applied to map the CIT168 probabilistic atlas labels to individual space (**Figure 1B**). Probabilistic label volumes were calculated by weighted summation of the label probability over the total volume using *fslstats (*FMRIB Software Library version 5.0.7). Probabilistic ROI volumes were calculated for the left and right hemisphere total amygdala and the following nine subregions: the lateral nucleus (LA), dorsal and intermediate divisions of the basolateral nucleus (BLDI), ventral division of the basolateral nucleus and paralaminar nucleus (BLVPL), basomedial nucleus (BM), central nucleus (CEN), cortical and medial nuclei (CMN), amygdala transition areas (ATA), amygdalostriatal transition area (ASTA), and anterior amygdala area (AAA). Finally, the amygdala RVFs for each subregion were calculated by dividing each region’s probabilistic volume by the total probabilistic volume of the hemispheric amygdala.

**Figure 1.**
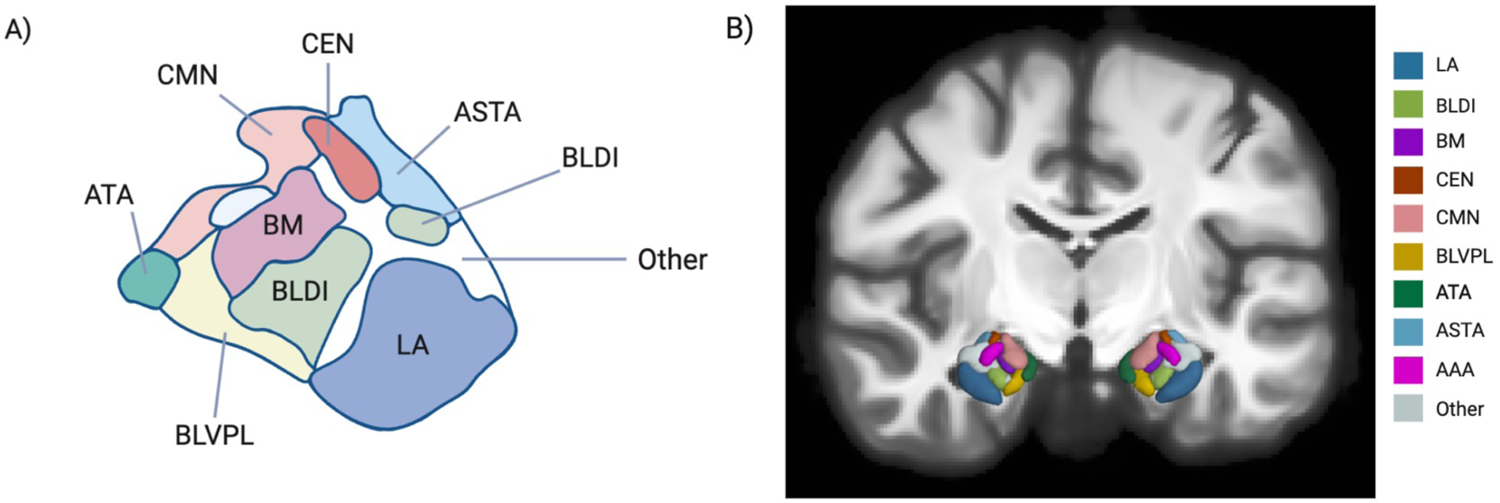
A) Schematic of CIT168 amygdala subregion labels. Note: The anterior amygdala area (AAA) is not visible in this view. Image created with BioRender. **B) Representative ABCD participant segmentations using the CIT168 Atlas.** 3D image created from in vivo segmentation of a representative participant using the Quantitative Imaging Toolbox (QIT) (76). Abbreviations: Lateral nucleus (LA), Basolateral Dorsal and Intermediate subdivision (BLDI), Basomedial nucleus (BM), Central nucleus (CEN), Cortical and Medial nuclei (CMN), Basolateral Ventral and Paralaminar subdivision (BLVPL), Amygdala Transition Area (ATA), Amygdalostriatal Transition Area (ASTA), Anterior Amygdala Area (AAA).

### 2.5 Demographics & Other Variables

Covariates included in our analyses were selected based on previous literature and the construction of a directed acyclic graph (**Figure S2**) (77). Sociodemographic variables such as race/ethnicity (*White, Black, Hispanic, Asian*, or *Other*), average household income in US Dollars (*<50K, 50-100K, >100K, Don’t Know/Refuse to Answer*), highest household education (*<High School Diploma, High School Diploma /GED, Some College, Bachelor Degree, Post-Graduate Degree, Missing/Refused*), urbanicity (*Rural, Urban Clusters, Urbanized*) and parent-report perceived neighborhood safety were included due to known disparities in air pollution exposure across socioeconomic status, race/ethnicity, and neighborhood safety, and known relationships between these variables and amygdala size (78–81). Measures of weekly physical activity (days per week) and average daily screen time (hours) were included, as previously published by our lab (82), since they may influence time outdoors, and thus exposure levels. We also included demographic factors and precision imaging variables such as the child’s age at scan, sex assigned at birth (*female* or *male*), body mass index (z-scored; BMIz), and handedness (*right, left,* or *mixed*), as well as MRI scanner headcoil (*32-* or *64-channel*) and MRI serial number, which account for site and scanner differences (70). Finally, intracranial volume (ICV) was included in models using absolute volumes to account for variation in total amygdala size.

### 2.6 Statistical Analyses

Analyses were conducted using *R* Version 4.3.2 (83). LME models were run to test the independent relationships between air pollution and total amygdala volume. That is, 36 single-pollutant models were run using the *lme4::lmer()* function to test associations between 18 outdoor air pollutants (three criteria pollutants,15 PM_2.5_ components) and left and right total amygdala volume. Models included the same covariates listed in **section 2.5** but implemented site as a random effect in place of MRI serial number. False discovery rate correction was implemented to correct for multiple comparisons (see **Supplemental Methods**).

PLSC analyses were conducted as previously reported (82,84,85). Briefly, PLSC is a multivariate statistical method that compares two multidimensional datasets that may have cross-correlated features (86) to identify patterns of shared covariance, or latent dimensions. To identify relationships between these datasets, PLSC employs singular value decomposition on the correlation matrix of these data matrices, identifying latent dimensions that capture the maximum shared covariance and the corresponding variables contributing to these latent dimensions, explained by their loadings. Considering the high multicollinearity between exposure to air pollutants and amygdala subregion volumes, PLSC is an optimal statistical approach due to its ability to handle highly cross-correlated data (see **Figures S3-5** for correlograms). We ran separate PLSC analyses to examine relationships between exposure to three criteria pollutants and 15 PM_2.5_ components and amygdala subregion volumes. Considering subregion sizes may vary as a function of total amygdala volume, we also conducted two complementary PLSC analyses using each set of exposures and relative proportions of amygdala subregions (i.e., RVFs). The PLSCs with probabilistic volumes were conducted to determine if air pollution exposure is linked to overall amygdala subregion volumes, whereas the PLSCs with RVFs were implemented to investigate if air pollution relates to differences in the relative subregion composition of the amygdala. Before execution of our PLSCs, five components (EC, NH_4_^+^, NO_3_^-^, OC, SO_4_^2-^) were converted from µg/m^3^ to ng/m^3^ so that PM_2.5_ components had identical units. To account for covariates (see **section 2.6.1**), we applied linear regression to residualize out the set of covariates from the data matrices, and then performed PLSC analysis on the residualized data. For permutation testing, data were resampled 10,000 times without replacement to identify significant latent dimensions. The probability of significance was determined based on the number of times permuted singular values exceeded the observed singular value, along with calculating the percentage of variance explained visually using scree plots. The bootstrap test evaluated the robustness of saliences loading onto significant latent dimensions by resampling the data 10,000 times, leaving out one sample each time. Confidence bootstrap ratios were derived by dividing the mean of a variable’s bootstrapped distribution by its standard deviation. Bootstrap ratios exceeding 2.5 (corresponding to p < 0.01) were considered statistically reliable and significant (87). The PLSC analysis was carried out using the *TExposition* and *TInPosition* packages (88), followed by a validation procedure with the *data4PCCAR::Boot4PLSC()* function (89).

## 3. Results

Sociodemographic and descriptive statistics for the final analytic sample (n = 4,473) are included in **Table 1** and **Tables S1** and **S2**. The final analytic sample had more male, Non-Hispanic White, and higher income individuals than the full ABCD sample, which is more urban, of higher socioeconomic status, and has higher levels of parental education than the U.S. population (90). Annual average exposures for the sample are presented in **Table S3**. Notably, PM_2.5_ exposure was 7.47 (1.47) µg/m^3^, NO_2_ was 19.3 (6.21) parts per billion (ppb), and O_3_ was 42.1 (4.47) ppb, which are significantly lower concentrations than the most recent EPA standards of 9 µg/m^3^ for PM_2.5_ (t(4,472) = −69.59, p < 0.001), 53 ppb for NO_2_ (t(4,472) = −363.19, p < 0.001), and 70 ppb 8-hour maximum for O_3_ (t(4,472) = −418.23, p < 0.001). These concentrations, however, are significantly higher than the World Health Organization (WHO) 2021 Air Quality Guidelines of 5 µg/m^3^ for PM_2.5_ (t(4,472) = 111.86, p < 0.001), 10 ppb for NO_2_ (t(4,472) = 100.14, p < 0.001), and significantly lower than the 60 ppb peak season 8-hour maximum for O_3_ (t(4,472) = −268.53, p < 0.001) (91). Descriptive statistics for total amygdala and subregion volumes and RVFs can be found in Table S4.

Using LME modeling, no significant associations were identified between pollutants and total hemispheric amygdala volumes (**Table S5**). Using PLSC, no significant latent dimensions were identified between exposure to criteria pollutants and amygdala subregion volumes (**Figure S6**). In contrast, one significant latent dimension was identified, explaining 52% of the shared variance, between exposure to the 15 PM_2.5_ components and amygdala subregion volumes (**Figure S7**). Specifically, higher Ca exposure was positively associated with volumes of the bilateral LA, BLDI, and BLVPL, as well as the left BM and AAA (Figure 2).

**Figure 2.**
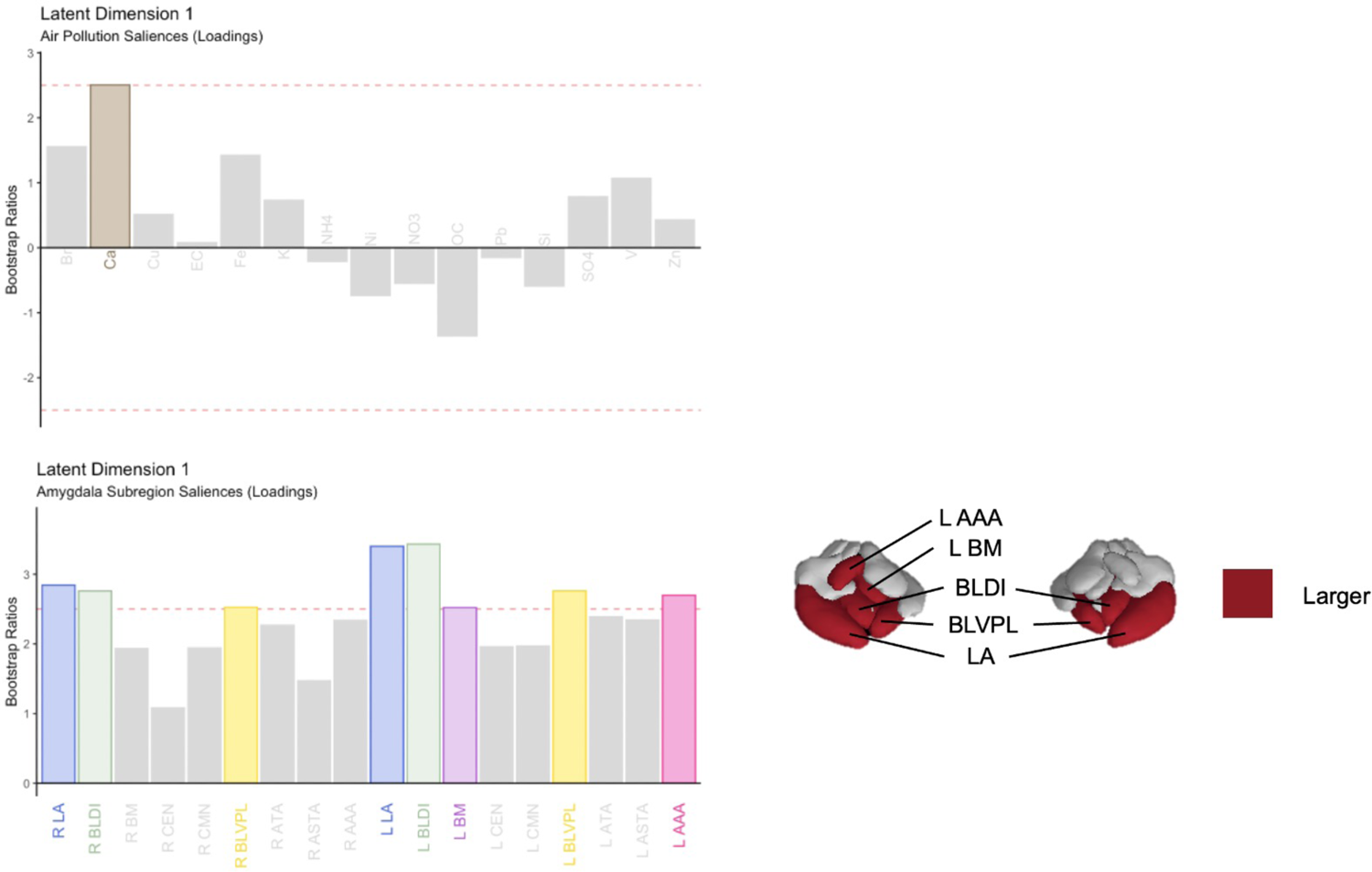
Variable loadings for the association between PM_2.5_ components and amygdala subregion volumes. Variables passing bootstrap ratio threshold (2.5; p < 0.01) are displayed in color. Significant subregions mapped into 3D brain space using in-vivo segmentation from a representative participant using the Quantitative Imaging Toolbox (QIT) (76); red denotes subregions with positive loadings (enlargement). Abbreviations: Bromine (Br), Calcium (Ca), Copper (Cu), Elemental Carbon (EC), Iron (Fe), Potassium (K), Ammonium (NH_4_^+^), Nickel (Ni), Nitrate (NO_3_^-^), Organic Carbon (OC), Lead (Pb), Silicon (Si), Sulfate (SO_4_^2-^), Vanadium (V), Zinc (Zn); Lateral nucleus (LA), Basolateral Dorsal and Intermediate subdivision (BLDI), Basomedial nucleus (BM), Central nucleus (CEN), Cortical and Medial nuclei (CMN), Basolateral Ventral and Paralaminar subdivision (BLVPL), Amygdala Transition Area (ATA), Amygdalostriatal Transition Area (ASTA), Anterior Amygdala Area (AAA).

PLSC analyses of amygdala apportionment revealed one significant latent dimension between criteria pollutants and amygdala subregion RVFs, explaining 82% of the variance (**Figure S8**). Within this significant dimension, PM_2.5_ was found to primarily drive these associations (Figure 3A). The PLSC between PM_2.5_ components and subregion RVFs identified two significant latent dimensions, explaining 39% and 30% of the variance, respectively (**Figure S9**). The first dimension suggested that overall co-exposure to the 15 PM_2.5_ components (i.e., no single pollutant driving the association based on variable loadings) was associated with increased RVFs of the bilateral LA and decreased RVFs of the bilateral BM, right CEN, and left CMN (Figure 3B). The second latent dimension was driven by higher exposure to K and OC relating to overall differences in RVFs (no specific region driving the association based on variable loadings) (Figure 3B).

**Figure 3.**
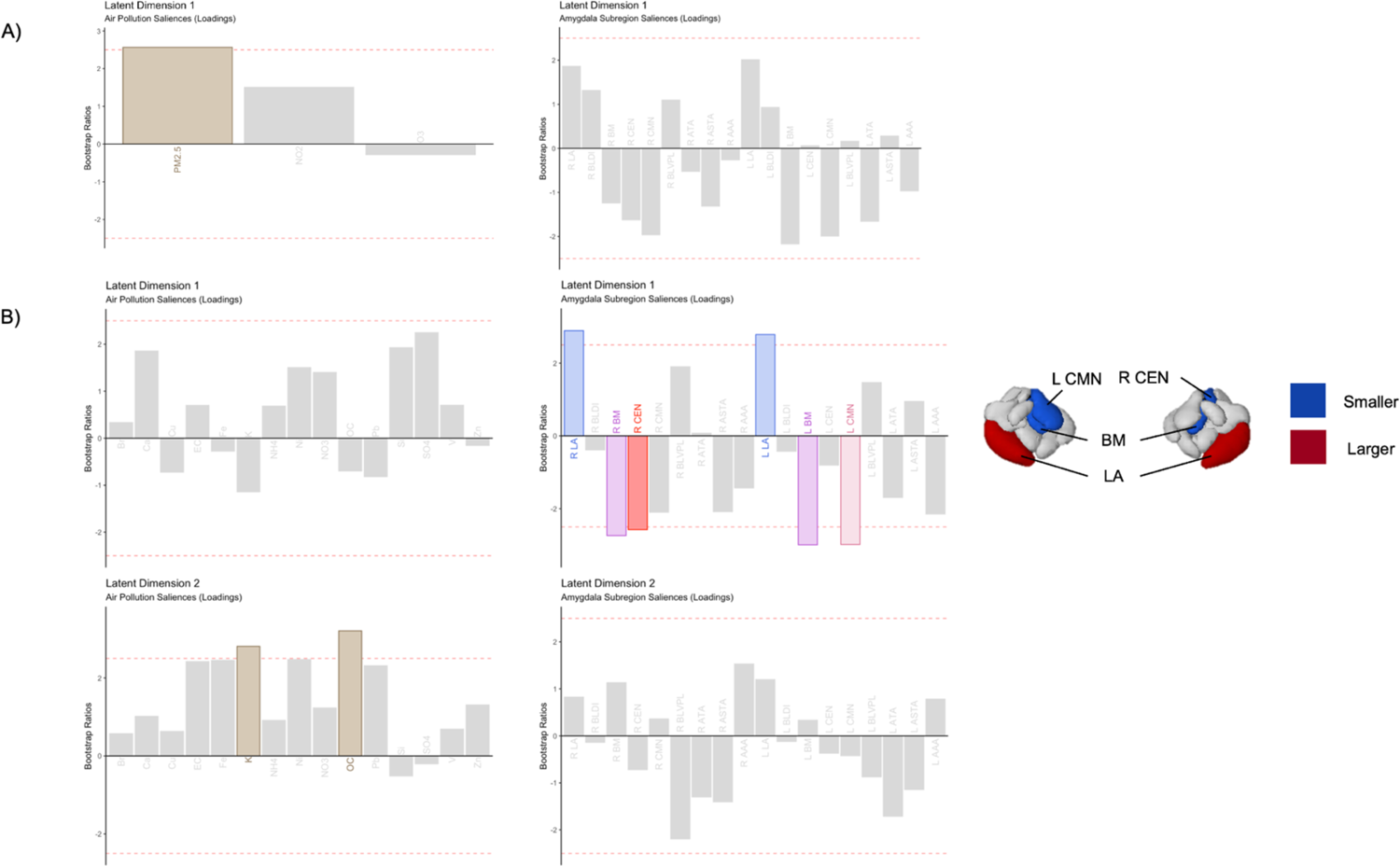
Variable loadings for criteria air pollutants, PM_2.5_ components, and amygdala subregion RVFs. A) Criteria Pollutants; B) PM_2.5_ components. Variables passing bootstrap threshold (p < 0.01) are displayed in color. Significant subregions mapped into 3D brain space for visualization purposes using the Quantitative Imaging Toolbox (QIT) (76); red denotes subregions with positive loadings (i.e. larger relative proportion with exposure) and blue denotes subregions with negative loadings (i.e., smaller relative proportion with exposure). Abbreviations: fine Particulate Matter (PM_2.5_), Nitrogen Dioxide NO_2_), ground-level ozone (O_3_); Bromine (Br), Calcium (Ca), Copper (Cu), Elemental Carbon (EC), Iron (Fe), Potassium (K), Ammonium (NH_4_^+^), Nickel (Ni), Nitrate (NO_3_^-^), Organic Carbon (OC), Lead (Pb), Silicon (Si), Sulfate (SO_4_^2-^), Vanadium (V), Zinc (Zn); Lateral nucleus (LA), Basolateral Dorsal and Intermediate subdivision (BLDI), Basomedial nucleus (BM), Central nucleus (CEN), Cortical and Medial nuclei (CMN), Basolateral Ventral and Paralaminar subdivision (BLVPL), Amygdala Transition Area (ATA), Amygdalostriatal Transition Area (ASTA), Anterior Amygdala Area (AAA).

## 4. Discussion

To our knowledge, this study is the first to explore the relationships between ambient air pollution exposure and amygdala subregion morphology. Previous studies have failed to identify consistent associations between childhood/adolescent exposure and total amygdala volumes (46,48–50), highlighting the importance of studying amygdala subregions. We aimed to identify novel associations between air pollution exposure and amygdala subregions by implementing a multivariate approach to better account for co-exposure to multiple pollutants in a large sample of preadolescents. Our results suggest that, while one year of exposure to any single pollutant does not relate to total amygdala volume, co-exposure to PM_2.5_ is associated with distinct amygdala subregion differences in early adolescence. Specifically, annual average PM_2.5_ and three PM_2.5_ components–Ca, K, and OC–are associated with distinct increases in basolateral volumes and differences in amygdala subregion apportionment at ages 9-10 years-old.

While total amygdala volumes increase through early adolescence (33,92), human postmortem (93) and neuroimaging studies (39,41,43) suggest heterogeneity in amygdala subregion development during childhood and adolescence. Previous studies have found nuclei specific changes in amygdala neuron numbers (93,94) and age-related differences in amygdala RVFs (61). Using these findings as context, the current study suggests that exposures to certain PM_2.5_ constituents may alter patterns of amygdala development. At 9-10 years, Ca was positively associated with several regions of the larger basolateral (BLA) complex (i.e., LA, BLDI, BLVPL, BM) and the AAA, which borders the BLA, separated from it by a thin band of fibers (52). Given that air pollutants were associated with distinct subregional volumes, but not total amygdala volumes, it is unsurprising that PM_2.5_ and its components were also related to patterns of regional apportionment. Specifically, exposure to PM_2.5_ components was associated with proportionally larger lateral, but smaller medial (i.e., BM and CMN) and central (CEN), subregions. A second association was noted between K and OC, attributes of biomass burning, and patterns of amygdala apportionment, albeit like total PM_2.5_ exposure, no single amygdala subregion drove these associations. Although this is the first study to examine air pollution and amygdala subregions, it supports a growing body of literature linking outdoor air pollutants with amygdala structure and function during development. In humans, higher prenatal exposure to coarse particulate matter and lower exposure to NO_2_ have been associated with smaller total amygdala volumes in infancy (45). Prenatal exposure to Si—a PM component in dust often coinciding with Ca—was related to larger total amygdala volumes in childhood, whereas prenatal polycyclic aromatic hydrocarbon and OC exposure were associated with smaller amygdala volumes at 9-12 years-old (46). Furthermore, in the ABCD cohort, our team previously identified associations between childhood PM_2.5_ exposure and longitudinal changes in resting-state functional connectivity of the amygdala and large-scale networks from ages 9 to 13 years-old (95), highlighting the impact outdoor air quality may have on the development of amygdala neurocircuitry.

Although additional research is needed, it is plausible that the observed relationships between air pollution exposure and amygdala subregions may have long-term implications for emotional processing and subsequent risk for mental health concerns. The BLA is the main thalamic sensory and cortical input region of the amygdala, involved in emotional regulation and processing and projecting high-level sensory input (96–98). The BLA plays a key role in conditioned fear and stress responses (96) and has been associated with an individual’s susceptibility to anxiety (99). The BM, which connects the LA and CEN, plays a significant role in the suppression of stress and fear responses, particularly in the context of social anxiety (100,101). The CEN receives intrinsic connections and is one of the major output nuclei of the amygdala; the CMN is another recipient of projections, particularly from the BLA, CEN, and olfactory bulb (52). Considering these roles in anxiety, fear conditioning, and social cognition (102), further investigations are needed to determine whether the identified subregion patterns play a role in the underlying neural mechanisms linking air pollution to risk for psychopathologies. An ongoing challenge, however, is the causal delay between the neurotoxicant effects of air pollution and observable behavioral differences. While several studies have identified positive associations between exposure to PM_2.5_ and anxiety and depression symptoms (14,15,103,104), some of them suggest a delayed onset between the timing of exposure and mental health concerns. For instance, a recent study showed that air pollution exposure at age 12 was not associated with concurrent mental health conditions, but rather higher incidence of depression at age 18 (105). As such, despite an absent association between annual PM_2.5_ and internalizing or externalizing symptoms in 9-13 year-olds in the ABCD Study (106), the current differences in the amygdala, alongside other notable outcomes (46,49,50,107–109), may reflect early biomarkers of neurotoxicity that ultimately contribute to increased risk for psychopathologies. Future longitudinal research is needed to determine how outdoor air pollution impacts trajectories of amygdala subregion development and apportionment and confirm its utility as a potential biomarker for later psychopathology.

Several strengths and limitations of the current study should be noted. We implemented the CIT168 atlas, which was created using in vivo Siemens MRI data from healthy young adult brains (51,52). While other amygdala segmentation approaches were created from post-mortem samples from older male brains (110,111), the CIT168 atlas uses high-resolution (700μm) Human Connectome Project data (from which the ABCD Study Siemens protocol was derived), along with probabilistic delineations to encode partial volume uncertainty in amygdala subregions. Moreover, to improve the reliability of our individual-level volume estimates, we chose *a priori* to limit our analyses to Siemens MRI data and use a stringent criterion for images based on CNR. While these strengthened the rigor of our amygdala subregion estimates, it limited our sample size to 4,473 participants from the larger ABCD Study. Our final sample included more male, White, and higher socioeconomic status participants, potentially limiting the generalizability of our findings. Furthermore, the participants included in this study are exposed to air pollution levels well below the EPA guidelines, though not below all WHO guidelines. While this study contributes to the growing knowledge base on associations between air pollution and the developing brain, these findings do not necessarily translate to adolescents living in highly polluted countries. Moreover, the data included in this study are cross-sectional; thus, while we can assess how current levels of air pollution relate to amygdala volume at one moment in time, we are unable to draw conclusions about the impact of chronic exposure on amygdala development.

To summarize, results of the current study suggest that exposure to PM_2.5_, specifically co-exposures to various components—calcium, potassium, and organic carbon—are related to differences in amygdala volumes and apportionment, with expansion in subregions involved in fear conditioning, and reduction in subregions responsible for anxiety and fear suppression. Taken together, these findings suggest air pollutant exposure may influence structural differences in amygdala subnuclei during a critical period of brain development. Future research investigating whether the current findings contribute to the biological underpinnings of the link between outdoor air pollution and risk for childhood and adolescent mental health conditions.

## Acknowledgments

A special thanks to the participants and families of the ABCD Study. Research described in this article was supported by the National Institutes of Health [MMH: NIEHS R01ES032295, R01ES031074; JMM and CCI: T32ES013678] and EPA grants [RD: 83587201, 83544101]. Data used in the preparation of this article were obtained from the Adolescent Brain Cognitive Development (ABCD) Study (https://abcdstudy.org), held in the NIMH Data Archive (NDA). This is a multisite, longitudinal study designed to recruit more than 10,000 children aged 9-10 and follow them over 10 years into early adulthood. The ABCD Study is supported by the National Institutes of Health Grants [U01DA041022, U01DA041028, U01DA041048, U01DA041089, U01DA041106, U01DA041117, U01DA041120, U01DA041134, U01DA041148, U01DA041156, U01DA041174, U24DA041123, U24DA041147]. A full list of supporters is available at https://abcdstudy.org/nih-collaborators. A listing of participating sites and a complete listing of the study investigators can be found at https://abcdstudy.org/principal-investigators.html. ABCD consortium investigators designed and implemented the study and/or provided data but did not necessarily participate in the analysis or writing of this report. This manuscript reflects the views of the authors and may not reflect the opinions or views of the NIH or ABCD consortium investigators. The ABCD data repository grows and changes over time. The ABCD data used in this report came from 10.15154/8873-zj65 and 10.15154/1520591. Additional support for this work was made possible from NIEHS R01-ES032295 and R01-ES031074. We would like to acknowledge Carinna Torgerson for her effort in processing the neuroimaging data used in analyses. We would also like to acknowledge Alethea de Jesus for data cleaning and preparation for analysis, and Jorge Max Landa and Jiawen Liang for assistance with creation of figure and table captions.

## CRediT Statement

Conceptualization: JM

Data Curation: JM, LNO, JS

Funding acquisition: MH, Formal Analysis: JM Methodology: JM

Project Administration: MH Resources: MH, JMT Software:

Supervision: MH Visualization: JM

Writing – original draft: JM

Writing – review & editing: MH, LNO, DLC, CCI, JS, JMT, JCC

