## Supplement for "Outdoor Air Pollution Relates to Amygdala Subregion Volume and Apportionment in Early Adolescents"

**Supplemental material for Outdoor Air Pollution Relates to Amygdala Subregion Volume and Apportionment in Early Adolescents**

Jessica Morrel, L. Nate Overholtzer, Kirthana Sukumaran, Devyn L. Cotter, Carlos Cardenas-Iniguez, J. Michael Tyszka, Joel Schwartz, Daniel A. Hackman, Jiu-Chiuan Chen, Megan M. Herting

**Supplemental Methods**

**Linear Mixed-Effect Models**

All statistical analysis was performed using *R* (Verson 4.3.2). In order to explore how air pollution impacts the whole amygdala, we ran 36 linear mixed-effects models using the *lme4::lmer()* function, testing the associations between each of the three criteria pollutants (fine particulate matter [PM_2.5_], nitrogen dioxide [NO_2_], ground-level ozone [O_3_]) and 15 PM_2.5_ components (Bromine [Br], Calcium [Ca], Copper [Cu], Elemental Carbon [EC], Iron [Fe], Potassium [K], Ammonium [NH_4_], Nickel [Ni], Nitrate [NO_3_], Organic Carbon [OC], Lead [Pb], Silicon [Si], Sulfate [SO_4_], Vanadium [V], Zinc [Zn]) and the total left and right hemisphere amygdala. Unlike partial least squares correlation (PLSC), which handles highly multicollinear data well, linear mixed-effect (LME) models require independent samples. Thus, to meet modeling assumptions, we had to randomly select 1 child per family, which reduced our analytic sample to 3,932 participants. Please see **Figure S1** for a flow chart of participant exclusion. Finally, because we are not able to include random effects in the residualization step of PLSC, we included MRI serial number as a proxy for site in those models. In our LME models, we replaced the fixed effect of MRI serial number with the random effect of site. The following formula was used:

*Amygdala Hemisphere Total Volume (L/R) ~ 1 + Air Pollutant/PM Component + Age + Sex + Race/Ethnicity + Household Income + Highest Parental Education + Physical Activity + Screen Time + Urbanicity + Neighborhood Safety + BMIz + Handedness + Headcoil + ICV (probabilistic analyses only) + (1 | Site)*

To correct for multiple comparisons, false discovery rate (FDR) correction was implemented using the Benjamini-Hochberg procedure and the *p.adjust* function in R. **Tables**

**Table S1. Analytic 3T Siemens MRI sample sites listed by geographic region.** Abbreviations: University of Colorado Boulder (CUB), Florida International University (FIU), Medical University of South Carolina (MUSC), Oregon Health & Science University (OHSU), University of Rochester (ROC), University of California - Los Angeles (UCLA), University of Florida (UFL), University of Maryland at Baltimore (UMB), University of Minnesota (UMN), University of Pittsburgh (UPMC), University of Utah (UTAH), Washington University in St. Louis (WUSTL), Yale University (YALE).

| **ABCD Site**  Northeast Region  ROC  UMB  UPMC  Yale  Southeast Region  FIU  MUSC  UFL  Midwest Region  UMN  WUSTL  West Region  CUB  OHSU  UCLA  UTAH | N = 4,473  238  338  257  284  398  231  274  382  384  352  378  272  685 |
| --- | --- |

**Table S2. Demographics table for final analytic sample, full ABCD sample, and responses from the 2016 American Community Survey (ACS)**. The total number of participants and (percentage of total sample) for each demographic category presented for the analytic sample, the total ABCD study sample, and a sample of US residents from the 2016 American Community Survey. Abbreviations: Adolescent Brain Cognitive Development (ABCD) Study; General Educational Development (GED), Body Mass Index (BMI). “Other” race/ethnicity category includes participants identified by their caregiver as American Indian/Native American, Alaska Native, Native Hawaiian, Guamanian, Samoan, Other Pacific Islander, Asian Indian, Chinese, Filipino, Japanese, Korean, Vietnamese, Other Asian not listed, or Other Race not listed. Associates degree and some college collapsed. ACS Urbanicity done on a decennial basis; ACS 2010 (Ratcliffe et al., n.d.)

|  | **Analytic Sample** | **Baseline ABCD Cohort** |
| --- | --- | --- |
| **Total Participants** | 4,473 | 11,839 |
| **Sex**  Female  Male | 1996 (44.6%)  2477 (55.4%) | 5658 (47.8%)  6181 (52.2%) |
| **Race/Ethnicity**  Asian  Non-Hispanic Black  Hispanic  Other  Non-Hispanic White  Missing/Refused | 62 (1.4%)  587 (13.1%)  807 (18.0%)  394 (8.8%)  2623 (58.6%)  0 | 250 (2.1%)  1777 (15.0%)  2405 (20.3%)  1243 (10.5%)  6162 (52.1%)  2 |
| **Household Income**  < $50k  ≥ $50k & < $100k  ≥ $100k  Don’t Know/Refused  Missing/Refused | 1067 (23.9%)  1241 (27.7%)  1847 (41.3%)  318 (7.1%)  0 | 3215 (27.2%)  3065 (25.9%)  4544 (38.4%)  1013 (8.6%)  2 |
| **Highest Parental Education**  < HS Diploma  HS Diploma/GED  Some College*  Bachelor Degree  Post-Graduate Degree  Missing/Refused  **Urbanicity**  Rural  Urban Clusters  Urbanized | 129 (2.9%)  390 (8.7%)  1111 (24.8%)  1256 (28.1%)  1581 (35.3%)  6 (0.1%)  335 (7.5%)  156 (3.5%)  3982 (89.0%) | 592 (5.0%)  1129 (9.5%)  3073 (26.0%)  3006 (25.4%)  4025 (34.0%)  14 (0.1%)  n = 11,186  964 (8.6%)  372 (3.3%)  9850 (88.1%) |
|  | **Mean (SD)** | **Mean (SD)** |
| Age (Months) | 119.7 (7.5) | 119.0 (7.5) |
| BMI Z-Score  Weekly Physical Activity (Days)  Avg. Daily Screen Time (Hours)  Neighborhood Safety | 0.3 (1.2)  3.6 (2.3)  3.0 (2.4)  3.9 (0.9) | 0.5 (1.2)  3.5 (2.3)  3.0 (2.4)  3.9 (0.9) |

**Table S3. Descriptive statistics and distributions of all criteria pollutants and fine particulate matter (PM_2.5_) components.** Abbreviations: nitrogen dioxide (NO_2_), ground-level ozone (O_3_); Bromine (Br), Calcium (Ca), Copper (Cu), Elemental Carbon (EC), Iron (Fe), Potassium (K), Ammonium (NH_4_), Nickel (Ni), Nitrate (NO_3_), Organic Carbon (OC), Lead (Pb), Silicon (Si), Sulfate (SO_4_), Vanadium (V), Zinc (Zn).

|  | N = 4,473  **Mean (SD)** | **Histogram** |
| --- | --- | --- |
| **Main Pollutants**  PM_2.5_ (*µg/m^3^*)  NO_2_ (*ppb*)  O_3_ (*ppb*) | 7.47 (1.47)  19.3 (6.21)  42.1 (4.47) | 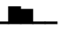  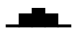  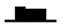 |
| **PM_2.5_ Components (*ng/m^3^*)**  Br  Ca  Cu  EC  Fe  K  NH_4_  Ni  NO_3_  OC  Pb  Si  SO_4_  V  Zn | 2.55 (0.54)  49.1 (21.1)  4.55 (1.57)  526 (139)  65.3 (21.7)  62.4 (9.10)  274 (116)  0.83 (0.25)  873 (334)  1828 (434)  4.55 (1.31)  88.7 (40.3)  875 (306)  0.39 (0.22)  9.12 (4.23) | 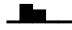  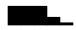  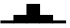  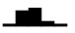  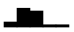  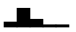  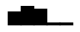  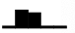  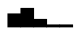  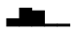  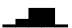  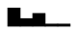  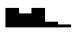  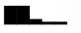  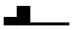 |

**Table S4. Descriptive statistics of probabilistic total amygdala and subregion volumes (mm^3^) and relative volume fractions (RVFs) by hemisphere in 9-10 year-olds from the ABCD cohort.** Abbreviations: lateral nucleus (LA), basolateral dorsal and intermediate subdivision (BLDI), basomedial nucleus (BM), central nucleus (CEN), cortical and medial nuclei (CMN), basolateral ventral and paralaminar subdivision (BLVPL), amygdala transition area (ATA), amygdalostriatal transition area (ASTA), anterior amygdala area (AAA), total hemispheric amygdala volume (total).

| **Right Hemisphere** | N = 4,473  **Mean (SD)** | **Left Hemisphere** | N = 4,473  **Mean (SD)** |
| --- | --- | --- | --- |
| LA  BLDI  BM  CEN  CMN  BLVPL  ATA  ASTA  AAA  Total | **Volume (mm^3^)**  325 (41.0)  188 (22.2)  105 (13.6)  46.6 (5.4)  158 (20.0)  116 (15.1)  83.7 (11.0)  70.0 (8.39)  57.8 (7.46)  1492 (159) | LA  BLDI  BM  CEN  CMN  BLVPL  ATA  ASTA  AAA  Total | **Volume (mm^3^)**  314 (39.5)  191 (22.6)  106 (13.8)  47.4 (5.52)  161 (20.7)  117 (16.2)  82.7 (11.2)  69.9 (8.44)  58.1 (7.58)  1483 (159) |
| LA  BLDI  BM  CEN  CMN  BLVPL  ATA  ASTA  AAA | **RVFs**  0.218 (0.014)  0.126 (0.004)  0.070 (0.004)  0.031 (0.002)  0.106 (0.007)  0.078 (0.005)  0.056 (0.004)  0.047 (0.004)  0.039 (0.003) | LA  BLDI  BM  CEN  CMN  BLVPL  ATA  ASTA  AAA | **RVFs**  0.212 (0.014)  0.129 (0.004)  0.072 (0.004)  0.032 (0.002)  0.108 (0.007)  0.079 (0.006)  0.056 (0.004)  0.047 (0.004)  0.039 (0.003) |

**Table S5. Associations between each air pollution A) left hemisphere and B) right hemisphere total amygdala volumes.** Results of 36 LME models run exploring associations between each of the three criteria pollutants and 15 fine particulate matter (PM_2.5_) components and the left and right hemisphere total amygdala volumes. Values reflect unstandardized beta coefficients, 95% confidence intervals (CI), and p-values (uncorrected and FDR corrected), adjusting for all confounders and covariates. Abbreviations: fine Particulate Matter (PM_2.5;_ *µg/m^3^*), Nitrogen dioxide (NO_2_; *ppb*), ground-level Ozone (O_3_; *ppb*); Bromine (Br), Calcium (Ca), Copper (Cu), Elemental Carbon (EC), Iron (Fe), Potassium (K), Ammonium (NH_4_), Nickel (Ni), Nitrate (NO_3_), Organic Carbon (OC), Lead (Pb), Silicon (Si), Sulfate (SO_4_), Vanadium (V), Zinc (Zn).

| **A) Total Left Hemisphere Amygdala** (N = 3,932) | | | | |
| --- | --- | --- | --- | --- |
|  | **beta (b)** | **95% CI** | ***Uncorrected p-value*** | ***FDR p-value*** |
| **Pollutant** |  |  |  |  |
| PM_2.5_ | 1.755430 | -1.014884 – 4.525744 | 0.214 | 0.718 |
| NO_2_ | 0.186978 | -0.525820 – 0.899775 | 0.607 | 0.743 |
| O_3_ | 0.221214 | -0.612803 – 1.055231 | 0.603 | 0.743 |
| Br | 6.152394 | -1.628376 – 13.933164 | 0.121 | 0.718 |
| Ca | 0.158119 | -0.079475 – 0.395714 | 0.192 | 0.718 |
| Cu | 2.056567 | -0.858288 – 4.971422 | 0.167 | 0.718 |
| EC | 0.015741 | -0.015966 – 0.047449 | 0.330 | 0.718 |
| Fe | 0.133547 | -0.077140 – 0.344234 | 0.214 | 0.718 |
| K | 0.180456 | -0.266012 – 0.626925 | 0.428 | 0.743 |
| NH_4_ | -0.011176 | -0.048860 – 0.026508 | 0.561 | 0.743 |
| Ni | 0.737851 | -15.896600 – 17.372301 | 0.931 | 0.976 |
| NO_3_ | -0.000207 | -0.013399 – 0.012985 | 0.976 | 0.976 |
| OC | -0.002556 | -0.012502 – 0.007389 | 0.614 | 0.743 |
| Pb | 0.866608 | -2.552966 – 4.286183 | 0.619 | 0.743 |
| Si | -0.062238 | -0.172995 – 0.048518 | 0.271 | 0.718 |
| SO_4_ | 0.001245 | -0.015617 – 0.018108 | 0.885 | 0.976 |
| V | 10.105673 | -9.551716 – 29.763062 | 0.314 | 0.718 |
| Zn | 0.478374 | -0.544315 – 1.501063 | 0.359 | 0.718 |
| **B)** **Total Right Hemisphere Amygdala** (N = 3,932) | | | | |
|  | **beta (b)** | **95% CI** | ***p-value*** | ***FDR p-value*** |
| **Pollutant** |  |  |  |  |
| PM_2.5_ | 1.771597 | -0.896825 – 4.440019 | 0.193 | 0.746 |
| NO_2_ | 0.140119 | -0.540636 – 0.820874 | 0.687 | 0.824 |
| O_3_ | -0.241117 | -1.061618 – 0.579385 | 0.565 | 0.765 |
| Br | 2.927375 | -4.361417 – 10.216166 | 0.431 | 0.746 |
| Ca | 0.103121 | -0.112197 – 0.318439 | 0.348 | 0.746 |
| Cu | 1.498422 | -1.336286 – 4.333130 | 0.300 | 0.746 |
| EC | 0.010880 | -0.019568 – 0.041329 | 0.484 | 0.746 |
| Fe | 0.077809 | -0.120388 – 0.276006 | 0.442 | 0.746 |
| K | 0.367251 | -0.047786 – 0.782288 | 0.083 | 0.746 |
| NH_4_ | -0.019985 | -0.054114 – 0.014144 | 0.251 | 0.746 |
| Ni | -4.357955 | -20.448617 – 11.732708 | 0.595 | 0.765 |
| NO_3_ | -0.001268 | -0.013630 – 0.011093 | 0.841 | 0.851 |
| OC | -0.000895 | -0.010252 – 0.008463 | 0.851 | 0.851 |
| Pb | 0.515931 | -2.757801 – 3.789663 | 0.757 | 0.851 |
| Si | -0.037617 | -0.146192 – 0.070957 | 0.497 | 0.746 |
| SO_4_ | -0.006894 | -0.021764 – 0.007977 | 0.363 | 0.746 |
| V | 9.362562 | -9.360133 – 28.085258 | 0.327 | 0.746 |
| Zn | 0.630101 | -0.334914 – 1.595115 | 0.201 | 0.746 |

**Figures**

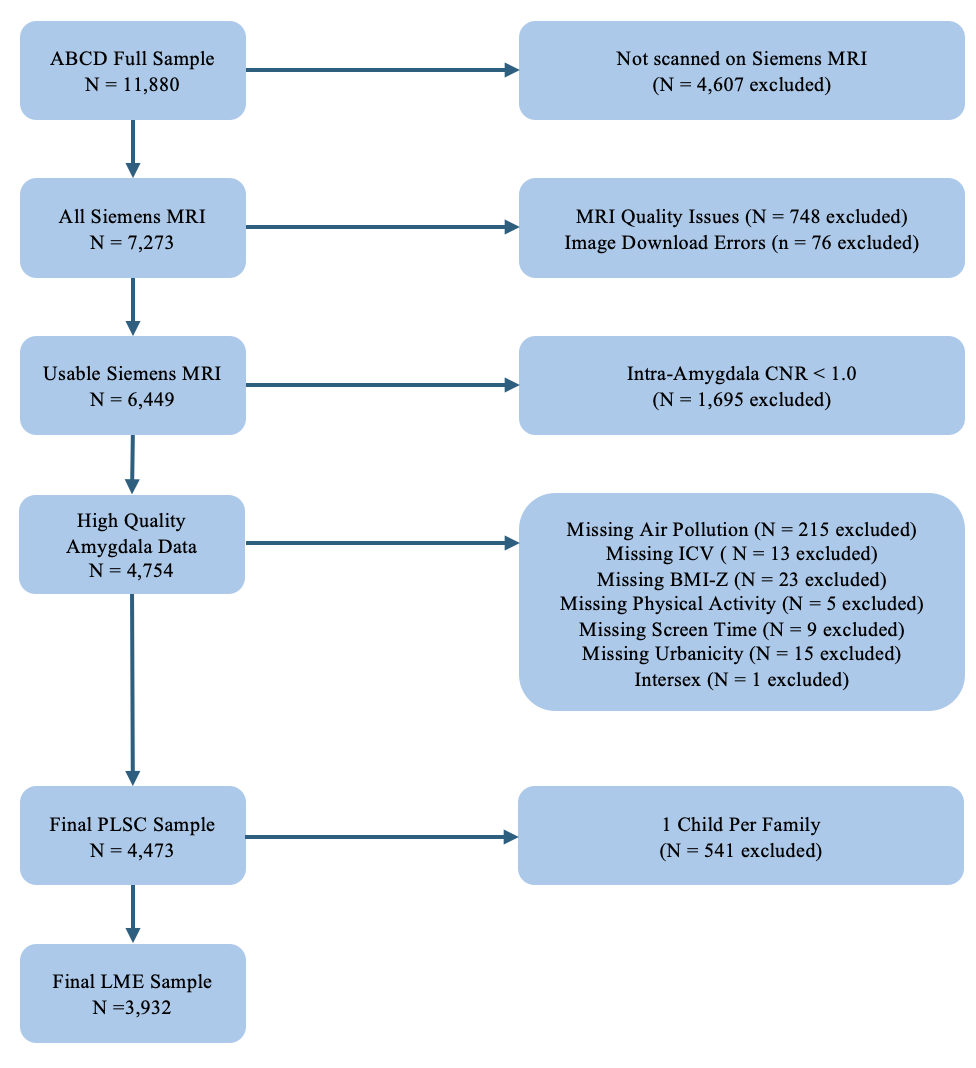

**Figure S1. Flowchart of exclusion and selection of final analytic sample.** Per PLSC requirements, the final sample has complete case-data for air pollution exposure, covariates, and high-quality brain outcome data. Abbreviations: Body Mass Index Z-score (BMI-Z), Contrast-to-Noise Ratio (CNR), Intracranial Volume (ICV), Linear Mixed-Effects Model (LME), Magnetic Resonance Imaging (MRI), Partial Least Squares Correlation (PLSC).

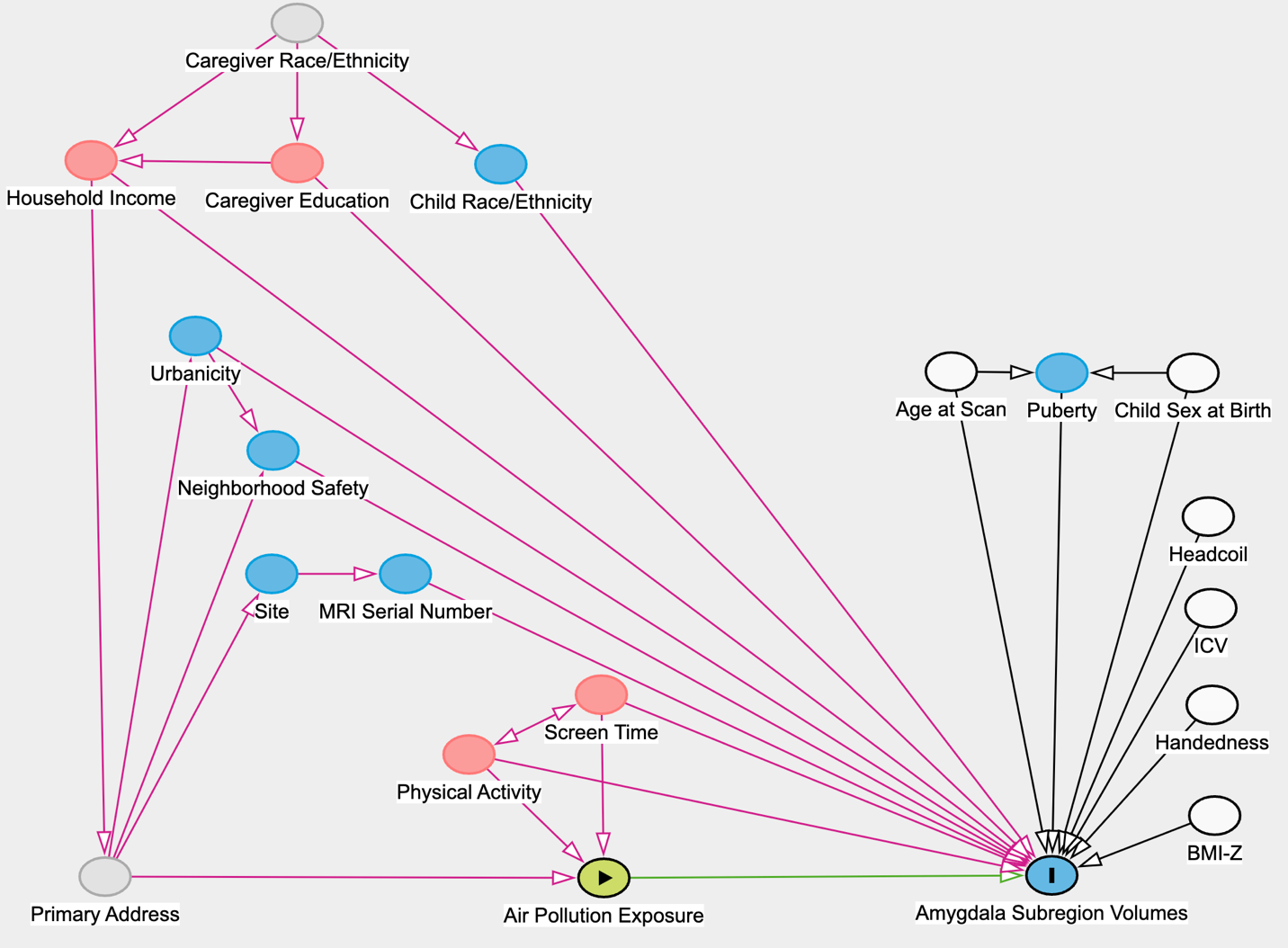

**Figure S2. Directed Acyclic Graph (DAG) displaying relationships between variables of no interest (i.e. covariates and confounders), air pollution exposure (predictor), and amygdala nuclei volumes (outcome).** Abbreviations: Body Mass Index Z-score (BMI-Z), Intracranial Volume (ICV), Magnetic Resonance Imaging (MRI).

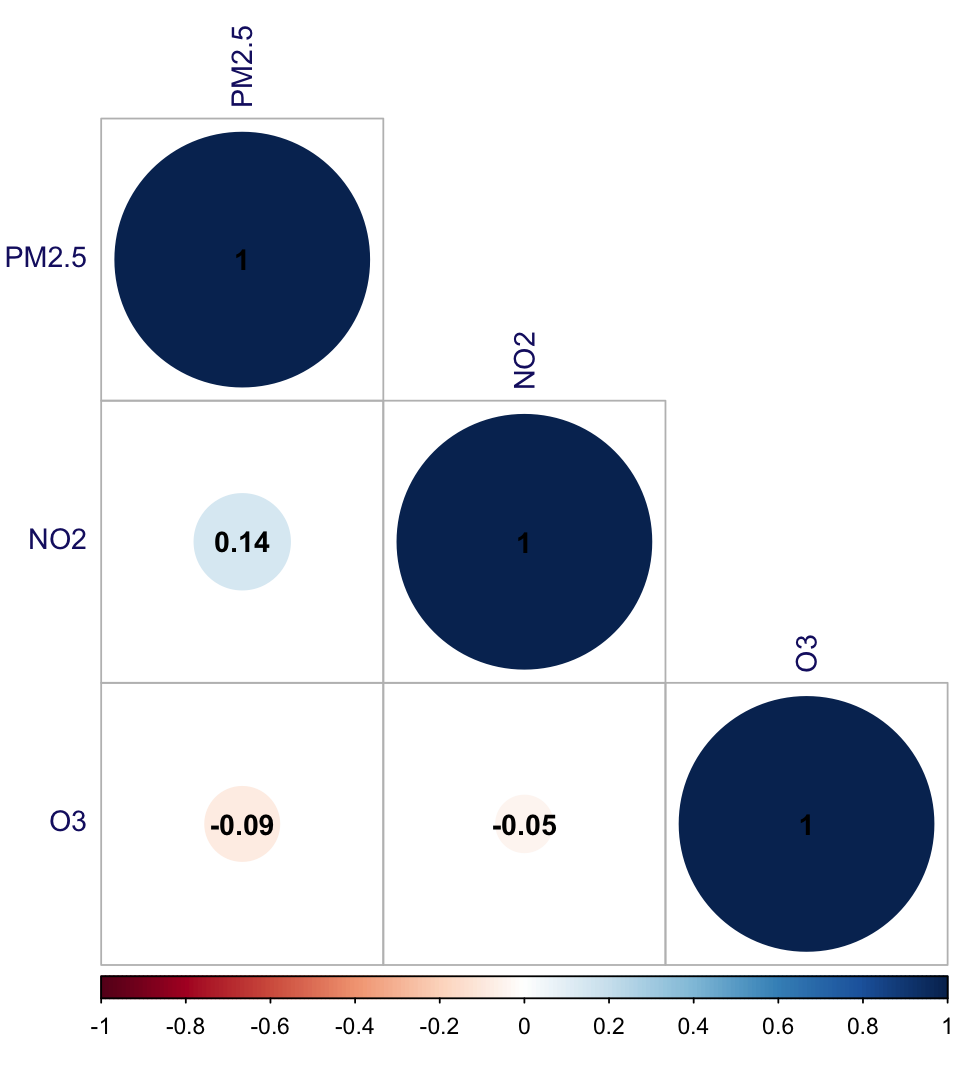

**Figure S3. Spearman’s correlation matrix of criteria pollutants.** Abbreviations: fine Particulate Matter (PM_2.5;_ *µg/m^3^*), Nitrogen Dioxide (NO_2_; *ppb*), ground-level Ozone (O_3_; *ppb*).

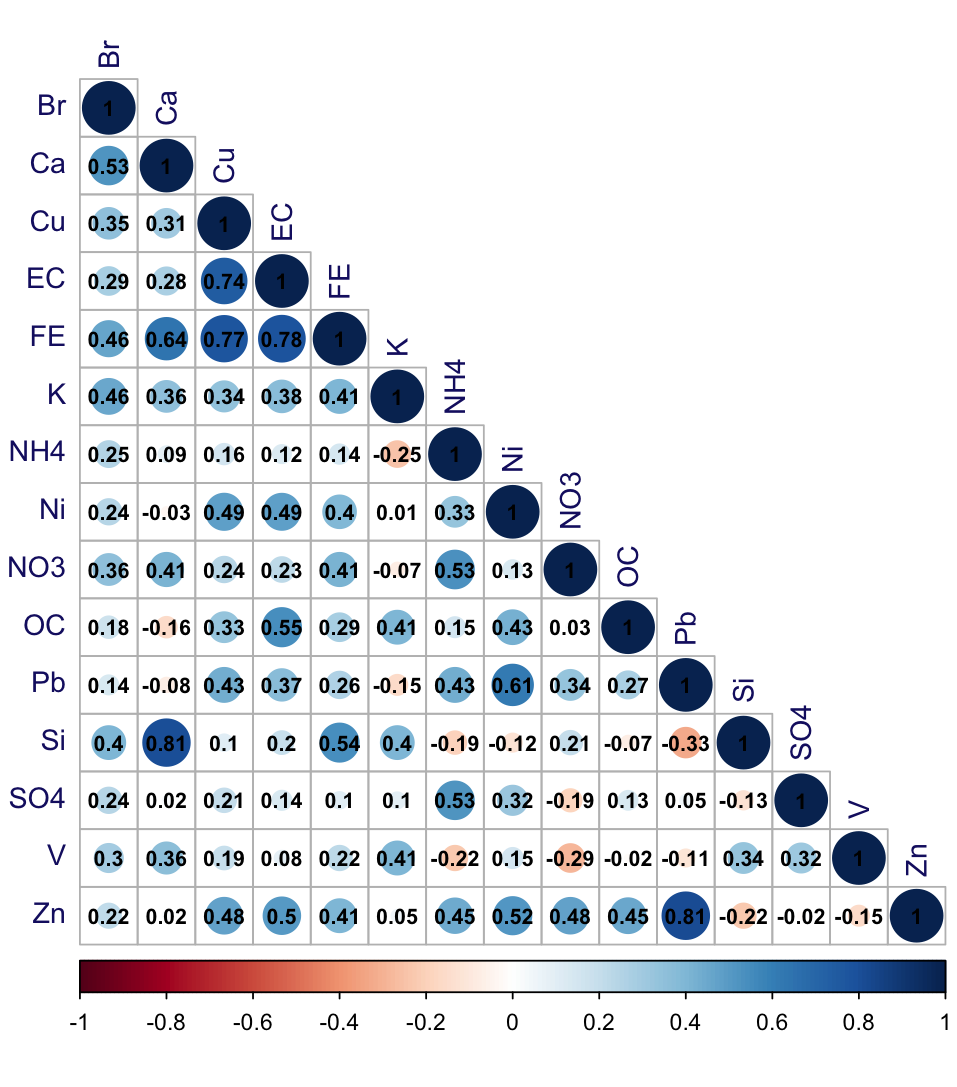

**Figure S4. Spearman’s correlation matrix of fine particulate matter (PM_2.5_) components (*ng/m^3^*).** Abbreviations: Bromine (Br), Calcium (Ca), Copper (Cu), Elemental Carbon (EC), Iron (Fe), Potassium (K), Ammonium (NH_4_), Nickel (Ni), Nitrate (NO_3_), Organic Carbon (OC), Lead (Pb), Silicon (Si), Sulfate (SO_4_), Vanadium (V), Zinc (Zn).

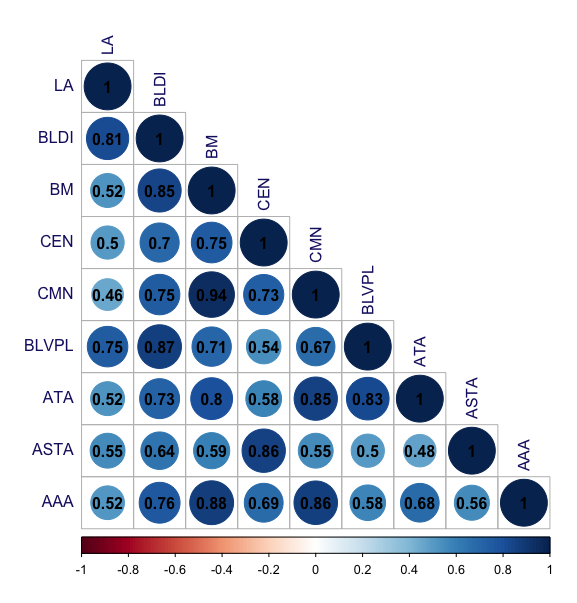

**Figure S5. Spearman’s correlation matrix of amygdala subregion probabilistic volumes (mm^3^).** Abbreviations: lateral nucleus (LA), basolateral dorsal and intermediate subdivision (BLDI), basomedial nucleus (BM), central nucleus (CEN), cortical and medial nuclei (CMN), basolateral ventral and paralaminar subdivision (BLVPL), amygdala transition area (ATA), amygdalostriatal transition area (ASTA), anterior amygdala area (AAA).

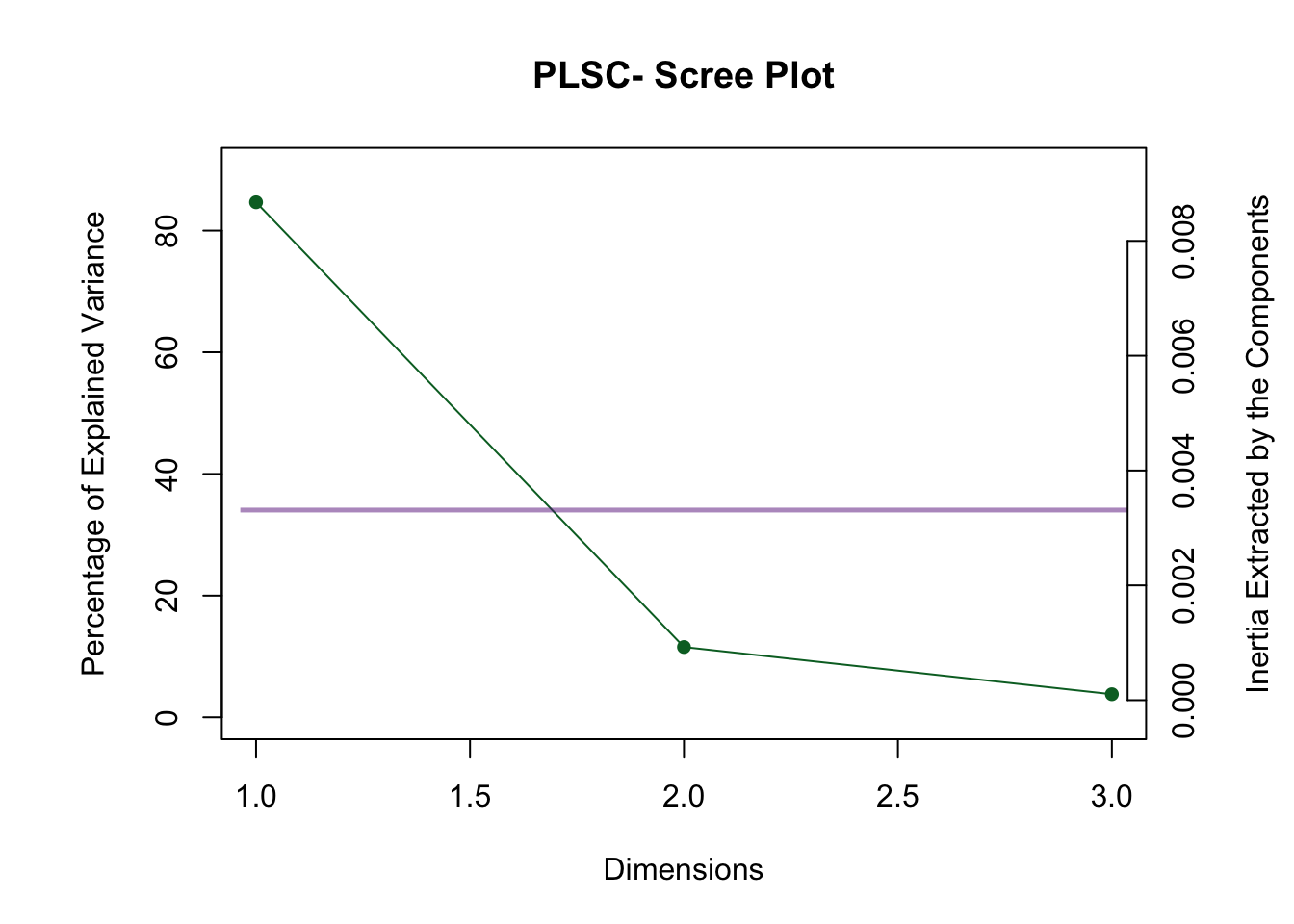

**Figure S6. Scree plot from the association between criteria air pollutants and amygdala probabilistic volumes**. Latent dimensions (x-axis) plotted alongside their associated percentage of explained variance (left y-axis) and inertia (right y-axis), which refers to each latent dimension’s sum of squared singular values. No violet bolded latent dimensions above the violet Kaiser criterion line indicates that there are no significant latent dimensions.

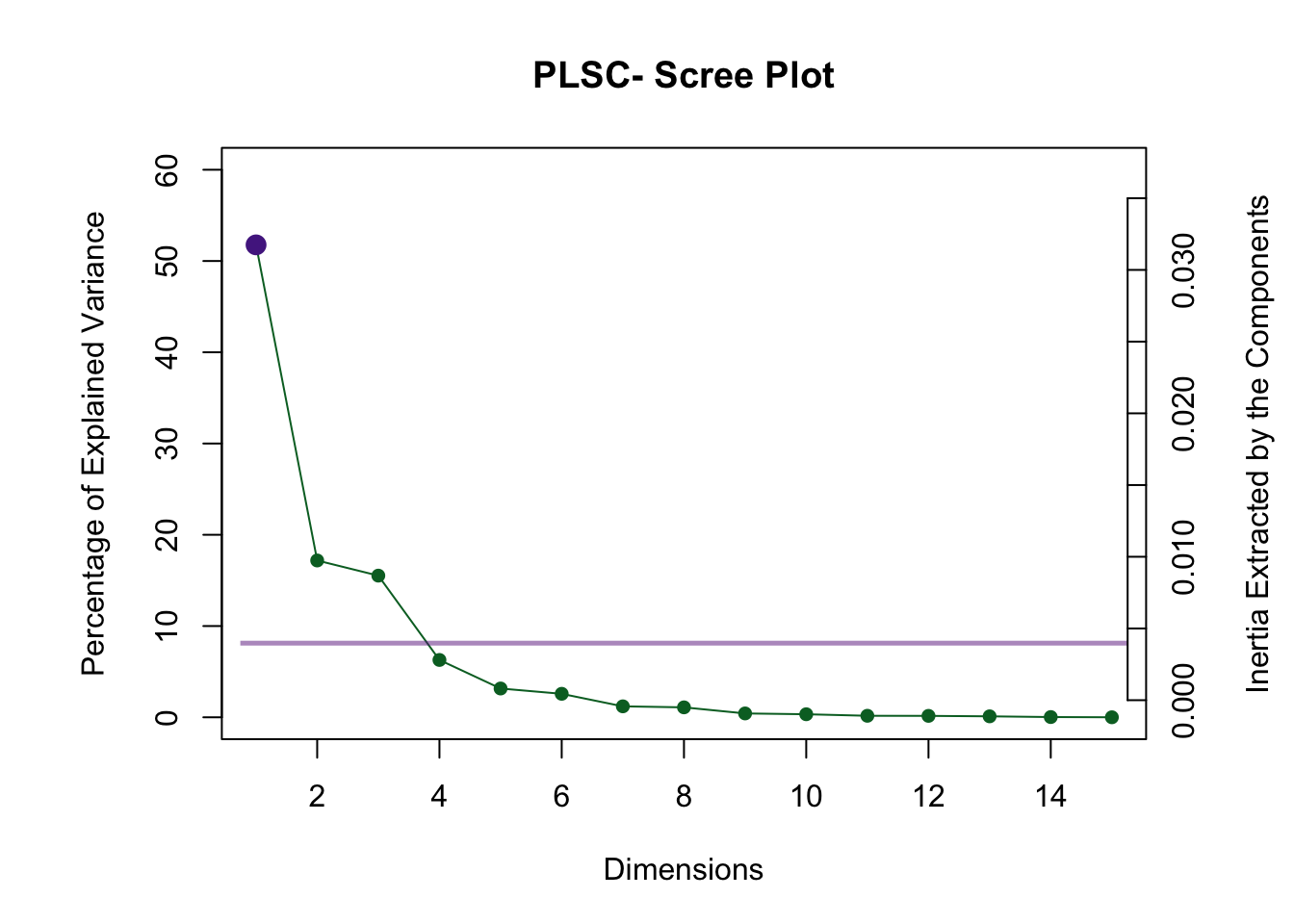

**Figure S7. Scree plot from the association between PM_2.5_ components and amygdala probabilistic volumes**. Latent dimensions (x-axis) plotted alongside their associated percentage of explained variance (left y-axis) and inertia (right y-axis), which refers to each latent dimension’s sum of squared singular values. The first violet bolded latent dimensions above the violet Kaiser criterion line represent the one significant latent dimension reported in results, explaining 52% of the variance.

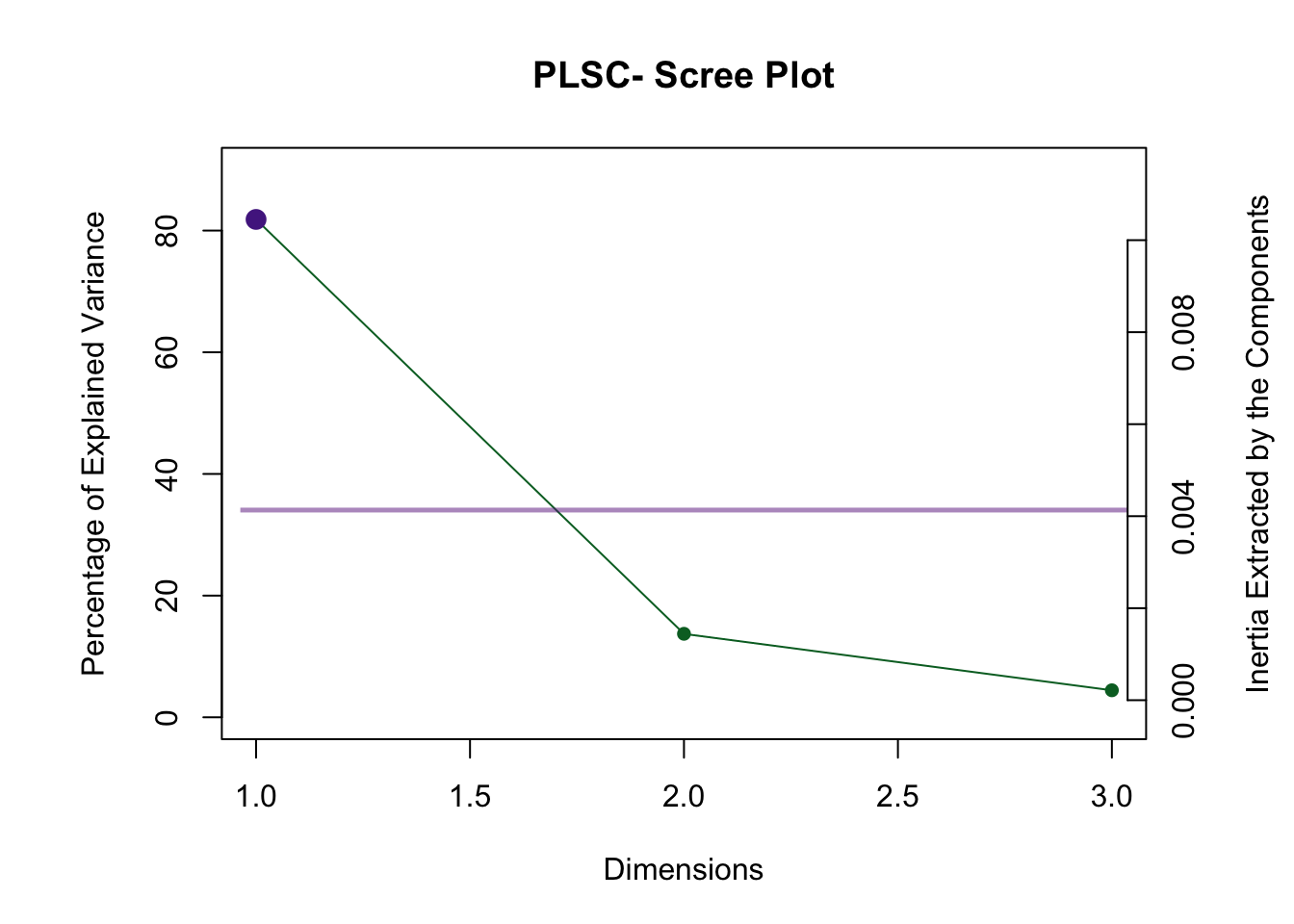

**Figure S8. Scree plot from the association between criteria air pollutants and amygdala relative volume fractions (RVFs)**. Latent dimensions (x-axis) plotted alongside their associated percentage of explained variance (left y-axis) and inertia (right y-axis), which refers to each latent dimension’s sum of squared singular values. The first violet bolded latent dimensions above the violet Kaiser criterion line represent the one significant latent dimension reported in results, explaining 82% of the variance.

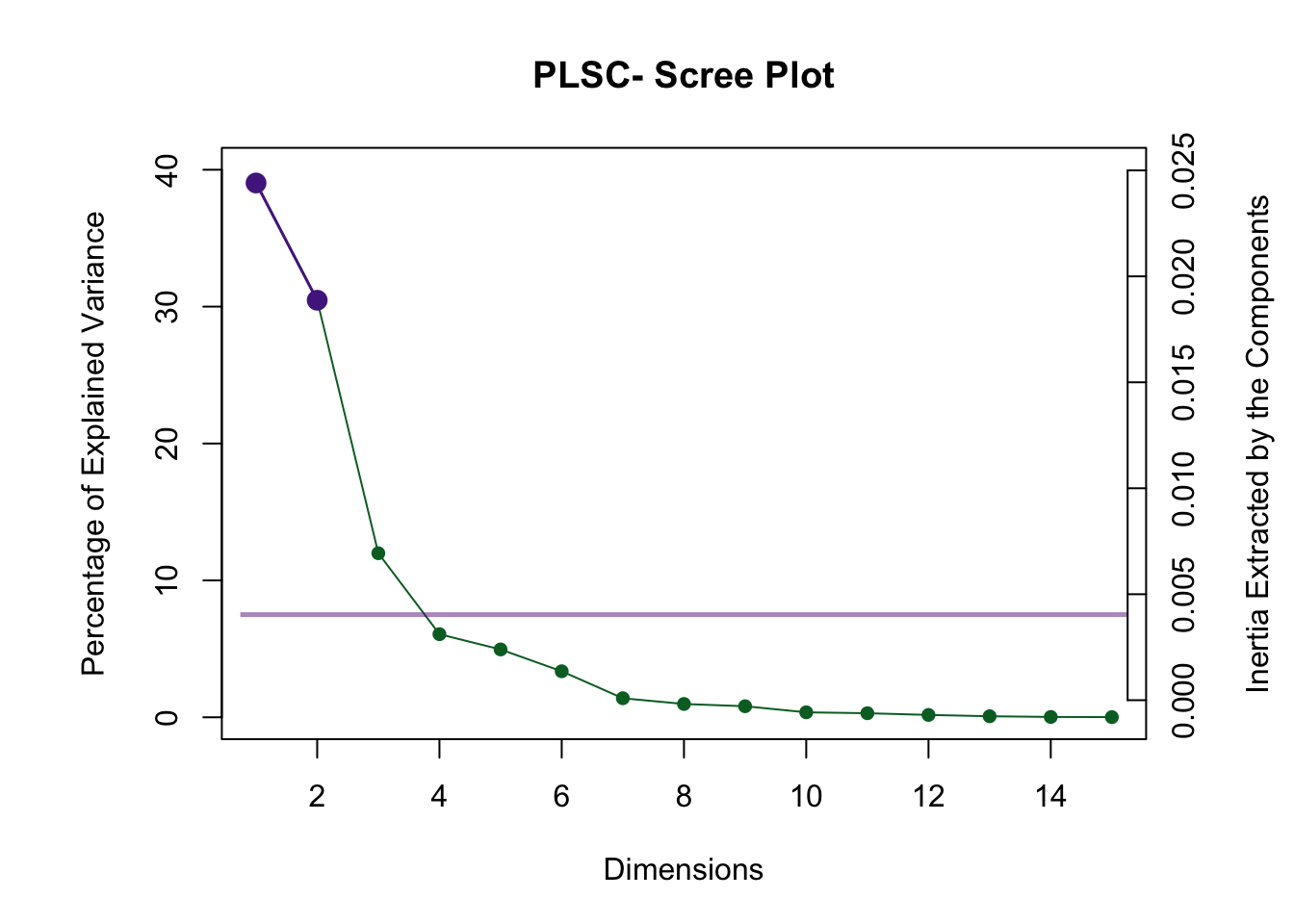

**Figure S9. Scree plot from the association between PM_2.5_ components and amygdala relative volume fractions (RVFs**). Latent dimensions (x-axis) plotted alongside their associated percentage of explained variance (left y-axis) and inertia (right y-axis), which refers to each latent dimension’s sum of squared singular values. The first two violet bolded latent dimensions above the violet Kaiser criterion line represent the two significant latent dimensions reported in results, explaining 39% and 30% of the variance respectively.
